# An Ethogram Identifies Behavioural Markers of Attention to Humans in European Herring Gulls (*Larus argentatus*)

**DOI:** 10.1101/2022.02.20.481240

**Authors:** Franziska Feist, Paul Graham

## Abstract

Herring gulls are one of few species thriving in anthropogenic landscapes. Their history of urbanisation and familiarity with people makes them a good target for studies of human-wildlife interactions. Previous research highlights a connection between food-stealing behaviour, success in anthropogenic areas, and increased attention towards humans, raising questions about the exact extent of a gull’s knowledge of human food cues. To explore these, behavioural responses to human cues in a food-related context were investigated and presented in a systematic ethogram, which identified three distinct markers of attention. Head turns, approaches, and angular body position all differed significantly between control and food conditions, showing that attention towards humans in a food-related context was upregulated and reflected in behaviour. In food condition trials, head turns occurred more often and gulls faced towards the experimenter with occasional approaches that were never seen in control conditions. Additionally, juvenile gulls exhibited less head turns, indicating lower vigilance, while neither approaches nor pecks differed between age groups. Interestingly, no group effects on attentional markers were found. Acoustic and behavioural human food-like cues alone seemed insufficient to elicit these responses, indicating that gulls specifically paid attention to the details of human behaviour. These results show situation-dependent attentional modulation in gulls and provide a description of attentive behaviours that can be used in further study.

**Summary:** Urbanised herring gulls successfully benefited from anthropogenic food sources. We showed that human food-centred behaviours directly modulated the attentional states of gulls.

## 1. Introduction

The human population is increasing, and with it comes the expansion of cities and urban landscapes (Lowry et al., 2013). Often this negatively affects local wildlife and causes species to avoid urbanised areas (Bateman and Fleming, 2012). Few manage to survive in anthropogenic landscapes, but those that do can be very successful and urban populations sometimes exceed densities found in natural environments (Bateman and Fleming, 2012).

This is true for the European herring gull (*Larus argentatus*). First instances of urban nesting occurred in the 1940s and by the 1960s, urban breeding colonies had spread throughout the United Kingdom’s coastal areas (Rock, 2005). Today, herring gulls’ overall population numbers are declining globally (BirdLife International, 2018), while urban populations continue to increase (Rock, 2005). As a result, urban herring gulls are familiar with the presence of people and anthropogenic food, which makes them an excellent target for studying the ecological and cognitive factors that contribute to the success of urban populations, but can also lead to human wildlife conflicts.

The kleptoparasitic behaviour of herring gulls suggests that they have a high degree of behavioural flexibility (Spencer et al., 2017) and an ability to store, integrate and use information about their environment and other individuals in decision-making (Morand-Ferron et al., 2007; Lee and Thornton, 2021), which may have been useful for adaptation to anthropogenic landscapes (Bateman and Fleming, 2012; Plumer et al., 2014). Urban gulls have been shown to pay attention to humans and human food cues, are aware of human gaze direction (Goumas et al., 2019), and preferentially peck at a food item that has been handled by a person when given the choice between two identical food objects (Kelley et al., 2020).

We investigated human-gull interactions in a food-related context by creating a systematic ethogram, a catalogue of behaviours exhibited by an animal, that described distinct attentive behaviours and confirmed whether urban gulls paid attention to humans in general or specifically in a food-related context.

## 2. Methods

Experiments were conducted by the same experimenter at the Brighton beachfront from March to April 2021 between 7:00 am and 11:00 am on weekdays only, which minimised the chances of pedestrian disturbances. No research was conducted on rainy days to protect the recording equipment. Pilot surveys indicated that gulls preferred to remain in the air on windy days (wind speeds > 22.5 km/h), and would not engage with the experimenter during low tide since they were preoccupied with natural foraging in tidal areas. Hence, no experiments were conducted on days on which low tide occurred between 7:00 am and 11:00 am. During all field research, government guidelines around Covid-19 were followed.

Recordings were made with a GoPro Hero 8 or an Honor 20 phone mounted on a LINKCOOL tripod. Video recording was paused or terminated when pedestrians walked into the camera’s field of view. It was always ensured that the animals were not disturbed or forced to engage with the experimenter. All gulls had the ability to remove themselves from the situation by either flying or walking away at any time during a trial. The experimenter adhered to the guidelines for the treatment of animals in behavioural research and testing by the Association for the Study of Animal Behaviour (ASAB) (Buchanan et al., 2012), and all necessary ethical approval was obtained from the University of Sussex.

### 2.1 Subjects

Single herring gulls (n=4) and groups (n=9) were approached for observations. For analysis, group sizes were split into Individual (n=4 birds), Small (≤ 10 birds in a group, n=4 groups with a total of 10 birds), Medium (11–20 birds, n=3 groups with a total of 33 birds), and Large (21+ birds, n=1 group with a total of 40 birds). The camera was set up about 10 m from the closest individual. We minimised chances for pseudo-replication by alternating trial locations, relying on the large local population size and this species’ fidelity to foraging areas (Clark, 2014; Fuirst et al., 2018; Lato et al., 2021).

Each group was observed for five to 20 minutes, depending on disturbances. Each five-minute block consisted of one of three conditions: a no-food condition (NFC) during which the birds were observed by the experimenter from behind the camera, a food condition (FC) during which the experimenter remained in the same position but retrieved a bag of red Walkers crisps from their backpack, pretending to eat from the bag, and a paper condition (PC) during which the experimenter took a piece of paper from their bag and simulated the rustling noises of a crisps bag by handling the paper along with eating and chewing motions. This would allow for a three-way comparison between a normal control (NFC), a food condition (FC) and a secondary control simulating acoustic and behavioural human food-like cues without presence of a real food item (PC). Other work (Feist et al., 2022 preprint) has shown that the colour of a crisp packet does not influence behavioural responses. If groups remained undisturbed and could undergo a complete recording session of 20 minutes, the following conditions were tested in random sequence: two NFCs, one FC, and one PC. The two NFCs resulted from the fact that the FC and PC conditions were split from each other by an NFC condition.

### 2.2 Video Analysis

Video files were analysed using the Behavioural Observation Research Interactive Software (BORIS) program (Friard and Gamba, 2016). General variables, such as weather condition (sun, cloudy, rain), average temperature in °C, and wind speed in mph during the time of recording, were noted along with low and high tide time and height on survey days. Later analysis showed that weather conditions on recording days only affected gull presence, but not their behaviour. All gulls were given a unique subject ID and their age was recorded as either juvenile, adult, or NA if unsure based on the recorded footage and the age classification provided in Mullarney et al. (2010). Recordings were then re-watched until the three selected behaviours of interest, head turns, body orientation relative to the experimenter, and approaches towards them, had been logged for all subjects; videos were rewatched so that on each viewing the experimenter could focus on a single gull. This effectively eliminated the chance of confusing birds within one video. The three selected distinct behavioural responses were hypothesised to be indicative of attention towards a person or situation (herein food).

### 2.3 Statistics and Modelling

To investigate whether the three selected behaviours were indeed markers of vigilance, statistical analysis focussed on comparing them across conditions. If they were indicative of an increase in attention in a food-related context, it was expected to observe more head turns and approaches during FC trials, and that gulls orient themselves more towards the experimenter. Gulls that participated in the experiment for less than 30 sec and those whose age could not be identified from the video footage (NA) were excluded from the analysis, resulting in a final n=194. Head turns and approaches were counted, and the weighted average angular body position (0–180°) of test subjects relative to the experimenter was calculated for each individual to obtain one measurement per bird per trial. Furthermore, head turns were divided by the time a gull was in camera view to give “head turns per gull minute”, which was used for analysis. All statistical analysis was done in R v.4.1.0 (R Core Team, 2021).

First, a generalized linear mixed model (GLMM) was constructed to investigate the effects of age (adults versus juvenile), condition (NFC, FC, and PC), and group size (range: 1–31 individuals; ordinal factor with four levels: individuals, small [2–10 gulls], medium [11–20 gulls], and large [21–31 gulls]) on head turns. Head turn data was log-transformed to improve the model diagnostics, and trial sequence (1–4) and date were included as random factors. To investigate the effects of age, condition, and group size on the angular body orientation while controlling for trial sequence and date, a second GLMM was built. The variance inflation factor (VIF) was used to investigate correlations among predictors for both models.

Lastly, Pearson’s Chi-square tests were conducted to confirm whether there was a relationship between either condition (NFC, FC, and PC), group size, or age and approach occurrence.

## 3. Results

The goal of this observational study was to create a systematic ethogram to identify markers of attention in a human food-related context. Our research suggests that attentive behaviours are upregulated during FC trials but not during NFC and only rarely during PC trials, which included simulated acoustic and behavioural food stimuli. Juveniles exhibited significantly less head turns than adults, while body orientation relative to the experimenter remained unaffected by age. Group size did not significantly affect head turns or body orientation relative to the experimenter.

### 3.1 The presence of a food stimulus increased head turn counts

The GLMM showed that head turns were significantly influenced by condition and age but not group size (*full vs. null:* x^2^ = 41.935, df = 6, p < 0.001). NFC differed significantly from FC (estimate = 0.351, se = 0.156, p < 0.05) but not from PC (estimate = −0.078, se = 0.190, p = 0.680). Additionally, there was a significant difference between FC and PC (estimate = −0.429, se = 0.209, p < 0.05). This confirms that head turns were upregulated in the presence of human food cues, suggesting increased vigilance in a food-related context (Fig. 1A). Additionally, juveniles showed fewer head turns compared to adults (estimate = −0.304, se = 0.131, p < 0.05), indicating that vigilance, as indicated by head turns, was lower in juveniles than in adults.

**Figure 1.**
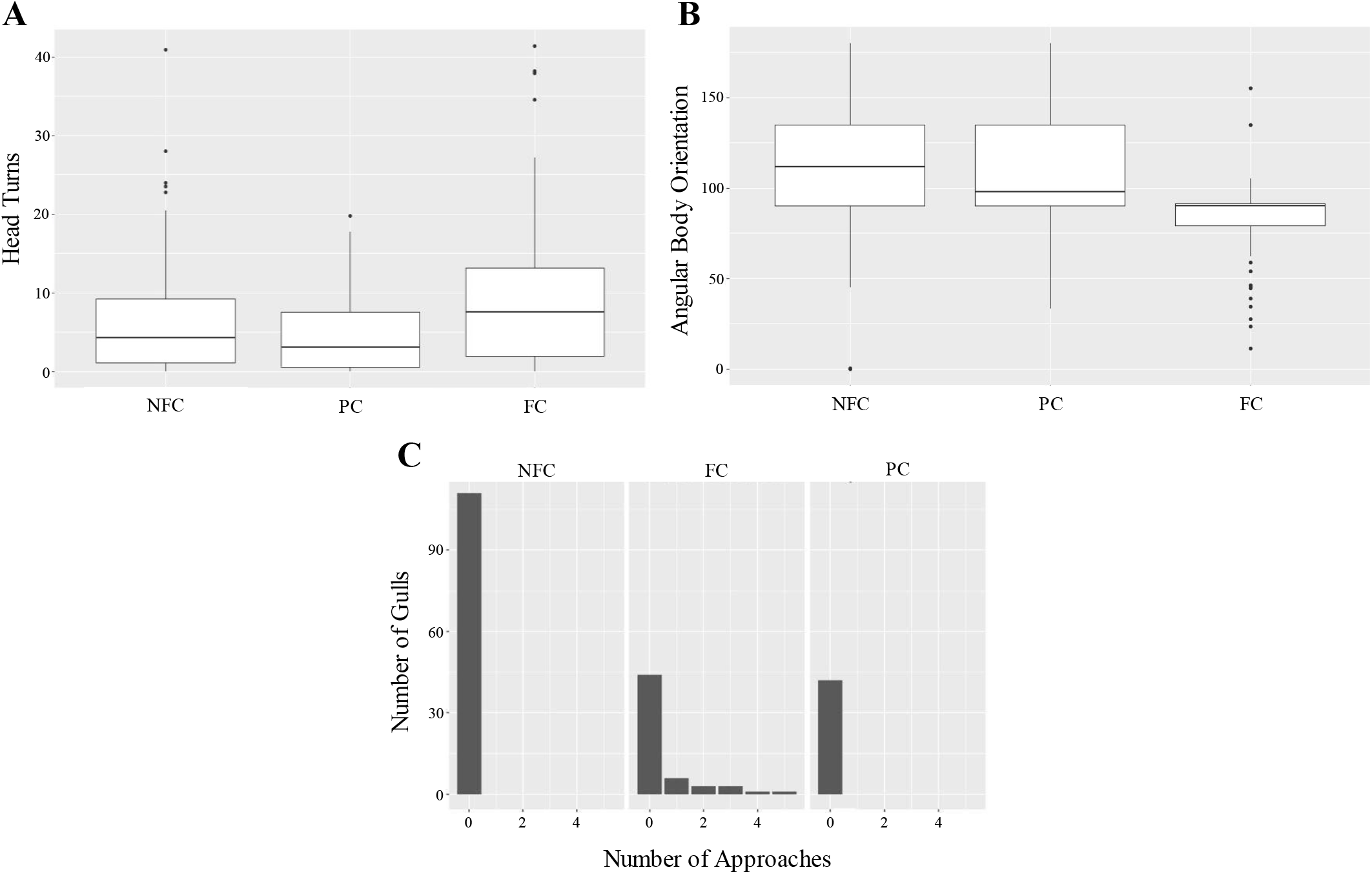
Upregulation of attentive behaviours during food trials. (A) The number of head turns of each subject per minute during each tested condition. During FC trials, head turns per minute were significantly higher than in NFC and PC trials (p < 0.05). The horizontal bars indicate the 1st and 3rd quartiles and the median. (B) The weighted average angular body position relative to the experimenter during the three conditions, with horizontal bars highlighting the 1st and 3rd quartiles and the median. During FC trials, orientations differed significantly from NFC trials (p < 0.001), with more gulls facing ≤ 90° towards and none facing > 115° away from the experimenter. (C) Approach instances during the three conditions. No gulls tested in either NFC or PC trials approached, while 14/53 or 26.41 % of gulls tested in FC trials did. Within one trial, gulls that did approach did so within one to five individual approach instances.

### 3.2 More gulls faced towards the experimenter in the food condition

The weighted average angular body position relative to the experimenter was significantly affected by condition (Fig. 1B), but not by age or group size (*full vs. null:* x2 = 41.671, df = 6, p < 0.001). Body orientation differed significantly between NFC and FC (estimate = −19.054, se = 5.120, p < 0.001) but not between PC and FC (estimate = −12.051, se = 6.334, p = 0.057) or NFC and PC (estimate = −7.003, se = 5.671, p = 0.217), with more subjects facing 90° or less towards the experimenter during FC trials.

### 3.3 Approaches only occurred when the experimenter signalled food

No trial effect on approach occurrence was found (Pearson’s Chi squared test, X-squared = 20.658, df =15, p = 0.148). No birds started approaching during NFC (n=101) or PC (n=40) trials, but 14 out of 53 gulls (26.42 %) did approach the experimenter during FC (n=53) trials, with between one and five approach instances per trial (Fig. 1C). A Pearson’s Chi squared test confirmed that there was an association between condition type and approach occurrence (X-squared = 41.142, df = 10, p < 0.001, n=194) but no significant effects of group size or age on approach occurrence.

Together, these results suggest that gulls paid increased attention to humans in possession of food (FC), which led to specific behavioural changes in food trials. Head turn counts increased significantly, more gulls oriented themselves towards or perpendicular to the experimenter, and some started approaching, which was not seen in other conditions. The acoustic and behavioural human food-like cues given during PC trials were not sufficient to elicit an increased number of head turns, a response only recorded in the presence of a real food item, and changes in body orientation were almost significant, with gulls facing slightly more towards the experimenter compared to NFC trials but less compared to FC trials.

## 4. Discussion

We have identified three behaviours: head turns, orientation towards an experimenter and approaches, that are upregulated during FC trials, which suggests that they are reflective of increased attention towards people in possession of food. It is hypothesised that all three behaviours allow gulls to combine attention towards a potential food source with higher vigilance and maintain possibility to escape if necessary. Since PC trials, during which human food-like behavioural and acoustic cues were mimicked, seemed insufficient to elicit head turn and approach responses similar to those seen in FC trials, they may be specific to a foraging context. Alternatively, the given cues may have been insufficient to signal food availability or deemed irrelevant by the gulls. The slight decrease in angle relative to the experimenter during PC trials may be indicative of some interest towards the paper, but the lack of approaches and head turns seems to highlight that gulls did differentiate between PC and FC contexts.

### 4.1 What purpose do these behaviours serve?

Approaches can be interpreted as interest in the experimenter, the food item and, potentially, a sign of preparations for a food-stealing attempt. They only ever occurred on trials where a food item was present, which supports this idea and confirms that the stimulus presented in this condition was attractive to the gulls. The approach rate of the studied population (24.14 %) was in line with what was expected based on previous reports and the experimental set-up used in this study. Previously, the approach rate of urban herring gulls towards a person in possession of food was reported to be around 26 % (Goumas et al., 2019). In contrast, approaches towards a food source 8 m away from the experimenter were around 68 % (Kelley et al., 2020). In our study, the food item remained with the experimenter and so the relatively low proportion of potential food-stealers (as indicated by approach attempts) is in line with our expectations. Head movements have previously been suggested to be a proxy for scanning behaviours (Fernández-Juricic, 2012) indicative of vigilance rates in birds (Hart and Lendrem, 1984). While foraging birds have been shown to be able to detect predation risks, raising their head for scanning increases their detection abilities (Lima and Bednekoff, 1999). Therefore, the increase in head turns can be seen as an indicator of increased vigilance towards an attractive food item. More frequent head movements may allow gulls to plan their approach accordingly, while simultaneously keeping watch for competition from conspecifics and predation risks.

In this regard, the lower number of head turns seen in juveniles may be indicative of juveniles’ general lower interest in an anthropogenic foraging opportunity (Monaghan, 1980), a lack of boldness required to engage with humans, or avoidance of competition with adults, in which they may be likely to lose.

Lastly, despite the non-significant result of the second GLMM, subjects were observed to face 90° or less towards the experimenter if a food object was present, which could be reflective of a gull’s attempt to prepare for an approach to steal food while ensuring that escape is possible in case the experimenter were to pose a threat. Predators preparing for an attack orient their head and body towards their target, which is used by prey to assess immediate predation risks (Book and Freeberg, 2015). At the same time, since humans can pose a threat to gulls (Goumas et al., 2020), they should orient themselves in a way that allows escape if necessary, such as facing predators side on, which increases predator detection (Kaby and Lind, 2003). Losing an encounter as prey will have more serious consequences than an unsuccessful foraging attempt (van den Hout et al., 2010), which may help explain why the birds tend to favour a position that trades escape opportunities with approach conditions. The fact that the majority of birds faced away from the experimenter during NFC trials may support this hypothesis, indicating that individuals may prioritise escape efficiency in situations where no foraging opportunity is available, or they may simply be disinterested. The orientations observed during PC trials were oriented more towards the experimenter than in NFC trials, but further away than in FC trials. This may indicate some interest in the handled paper, albeit less than in the real food items presented in FC trials.

### 4.2 How do gulls identify food objects?

While we did show that the specific food stimulus we presented during FC trials seemed to be attractive to the gulls, this interpretation does open up questions about the mechanisms of anthropogenic food choice in gulls. Our PC control trials, which consisted of human behavioural and acoustic food-like cues without an actual food item, were seemingly insufficient to reliably signal food availability, potentially owing to the fact that the given cues were not considered relevant enough. Both this and previous studies highlight the fact that gulls specifically pay attention to humans in possession of food, and that these combined cues influence approach (Goumas et al., 2019; Kelley et al., 2020). Interestingly, when given the choice between two non-food items, gulls do not peck at a handled non-food item above chance levels (Kelley et al., 2020), indicating that gulls may make a decision about the nature of an object based on the object’s characteristics before attending to human cues associated with it. As such, the object itself may be used to determine whether it is indeed a food item, while human cues are used to make choices about foraging opportunities. This notion would explain why the behavioural and acoustic cues given in the PC trials were insufficient to elicit an increase in head turns similar to that seen when a real food item was presented; gulls may have identified the object as a non-food object without the need for human cues and thus paid less attention to the situation.

In contrast, the lack of increased head turns or approaches in PC trials may be due to the behavioural and acoustic cues having been insufficient for gulls to identify a foraging opportunity. Little is known about what types of sensory cues gulls use or need to identify food items, but the topic of food choice has been studied in a variety of other species. In primates, visual cues are used most often to identify familiar food items, while the identification of novel items relies on olfactory, gustatory, visual and tactile cues (Laska et al., 2007). Interestingly, the importance of each type of sensory cue varies between species (Laska et al., 2007) and is associated with their foraging ecology (Rushmore et al., 2012). To reliably identify preferred food items, folivores require both visual and olfactory cues, while generalists could identify their foods using either cue alone, and frugivores use olfactory cues alone (Rushmore et al., 2012). The importance of olfactory cues in food choice is further highlighted by studies of a variety of taxa: fruit bats (*Cynopterus sphinx*) (Acharya et al., 1998); Loggerhead Sea turtles (*Caretta caretta*) (Pfaller et al., 2020); and sea birds (African penguins (*Spheniscus demersus*): Cunningham et al., 2008; Cory’s shearwater (*Calonectris borealis):* Bastos et al., 2020). Additionally, a variety of seabird species rely on olfaction for successful migration (lesser black-backed gulls (*Larus fuscus fuscus*): Wikelski, 2015), individual recognition (king penguins (*Aptenodytes patagonicus*): Cunningham and Bonadonna, 2015), and other tasks (common diving petrels (*Pelecanoides urinatrix Gmelin*) and thin-billed prions (*Pachyptila belcheri Matthews*): Bonadonna et al., 2003) (Nevitt, 2008; Schreiber and Burger, 2001). As such, this sense may also be used by herring gulls for tasks such as identification of foraging opportunities, which could explain their lowered attention in a context that lacked olfactory cues (PC).

### 4.3 Application of the presented ethogram

Our results indicated that urban herring gulls modify their behaviour in response to humans when food is present. Head turns, approaches, and body orientation relative to the experimenter were upregulated only when gulls paid attention to a person in possession of food. As such, these behaviours can be used as markers of attention towards human cues, which will, in turn, be useful in future studies of herring gull behaviour and attention. It has been suggested that gulls’ success in urban environments may be due to their cognitive capabilities and high behavioural flexibility (Bateman and Fleming, 2012; Plumer et al., 2014), and adaptive modulation of attention may play a role in this. One important unanswered question is how food-dependent modulation of attention develops in urban herring gulls. We only observed a significant difference in head turns exhibited by adults and juveniles; however, herring gulls have a juvenile period of multiple years (Mullarney et al., 2010) and differences between first winter juveniles and older individuals may be present. More detailed investigations of potential age differences, e.g., younger compared with older juveniles, would highlight whether this specific attentional modulation is a learned skill and, if so, when it develops.

Similarly, we may want to ask whether a gull’s response would be the same when they pay attention to other animals they have identified as a potential target for kleptoparasitism, or whether these behaviours are human-specific. Previous studies have shown that gulls specifically pick more successful individuals (i.e., those in possession of a larger food item) as food-stealing targets (Busniuk et al., 2020), which may imply that knowledge about their targets allows them to modulate their approach strategy accordingly. As a result, their behavioural response to a non-human individual in possession of food may differ from the responses we present here. Comparisons of how urban gulls attend to food cues from other species and how non-urban gulls react to human food cues will provide more detailed insight into whether urbanisation and frequent contact with people may have caused specific human-centred behaviours to arise in urban populations.

Lastly, with the ethogram presented here it will be possible to investigate whether attentional cues can be transferred across a group, even when direct visual cues of the food stimulus are lacking. Previous investigations of vigilance in herring gull groups have reported mixed results. Some highlight that vigilance spreads throughout a group, with individuals interrupting their sleep more often to scan their environment if their neighbours are more vigilant (Beauchamp, 2009). Others suggest that groups should follow the “many eyes” hypothesis, according to which individuals decrease their own vigilance when surrounded by highly vigilant neighbours (Roberts, 1996). Using the above identified attentional markers we can investigate how vigilance travels through a group of urban gulls, and how the presence of human food cues could affect this.

## Data availability

The data are available upon request.

## Competing interests

We declare we have no competing interests.

## Acknowledgements

We thank Professor Pierre Nouvellet (University of Sussex) for his support with the statistical modelling of the data.

## Funding

Not applicable.

## Notes

### Competing Interest Statement

The authors have declared no competing interest.

### Summary of Updates

Models were updated to investigate age and group effects on behavioural markers.

